# Effect of different ambient temperatures on reproductive outcome and wellbeing of lactating females in two mouse strains

**DOI:** 10.1101/2021.10.15.464536

**Authors:** Thomas Kolbe, Caroline Lassnig, Andrea Poelzl, Rupert Palme, Kerstin E. Auer, Thomas Rülicke

**Author notes:** Corresponding author (TK).

## Abstract

Ambient temperature is an important non-biotic environmental factor influencing immunological and oncological parameters in laboratory mice. It is under discussion which temperature is more appropriate and whether the commonly used room temperature in rodent facilities of about 21°C represents a chronic cold stress or the 30°C of the thermoneutral zone constitutes heat stress for the animals. In this study we selected the physiological challenging period of lactation to investigate the influence of a cage temperature of 20°C, 25°C, and 30°C, respectively, on reproductive performance and stress hormone levels in two frequently used mouse strains. We found that more pups were weaned from B6D2F1 hybrids compared to C57BL/6N mothers and that the number of weaned pups was strongly reduced if mothers of both strains were kept at 30°C. Furthermore, at 30°C mothers and pups showed reduced body weight at weaning and offspring had longer tails. Despite pronounced temperature effects on reproductive parameters, we did not find any impact on adrenocortical activity in breeding and control mice. Independent of the ambient temperature however, we found that females raising pups showed elevated levels of fecal corticosterone metabolites (FCMs) compared to controls. Increased levels of stress hormone metabolites were measured specially around birth and during the third week of lactation. Our results provide no evidence for reduced or improved wellbeing of lactating mice at different ambient temperatures, but we found that a 30°C cage temperature impairs reproductive performance.

## Introduction

Aiming to study thermoregulatory behavior in mice Gordon and Coworkers [1] started a discussion about the optimal ambient temperature, which culminated in a widely noticed publication of Hylander and Repasky [2]. The authors emphasized in their paper the different results of immunological and oncological studies when conducted at 20°C or at 30°C. Consequently, the results of studies on mouse models for human diseases, performed at 20-26°C standard ambient temperature were questioned and considered to be temperature biased, because of low reproducibility if performed under higher ambient temperatures [3-6]. It is generally accepted that room temperature can influence experimental results, like many other biotic and non-biotic environmental factors [7]. However, some of the reported effects related to ambient temperature merge only when mice were heated up to a body temperature of 39-40°C for 6 h [8-12] or to 42°C for 40 min [13].

Although a comprehensive analysis about the appropriate ambient temperature for laboratory mice in experiments is still missing, the call for housing laboratory mice in their thermoneutral zone as standard ambient temperature arised. The thermoneutral zone is defined as a temperature range in which the general metabolism of the organism, in the absence of any physical activity, generates sufficient heat as a byproduct of the continually ongoing metabolism to maintain the predetermined body temperature [14]. Thermal physiology of nocturnal mice seems to be different between dark and light periods. Influenced by the circadian rhythm two diurnal changing discrete ambient temperatures are proposed as thermoneutral points (TNP): ∼29°C in light phase and ∼33°C in dark phase [15]. In initial tests mice preferred to stay in warmer areas of experimental settings even if nesting material was provided. These thermoregulatory experiments were conducted using a copper pipe with a wire mesh inside [1] or an aluminium channel [16], heated at one side, cooled at the opposite side. This setup led to the assumption that mice prefer an ambient temperature near their homeothermic temperature of 30°C. In later studies, a more common laboratory mouse environment was used [17,18]. By offering bedding and nesting material it became obvious that the preferred ambient temperature depends on the activity of the mice and the amount and quality of nesting material [19-23]. With enough and useful nesting material mice can prevent their body from cooling down during resting periods [24]. Depending on activity, the body core temperature can change between 36°C and 37°C [25]. Also, the homeothermic zone seems to be more a temperature point than a zone and varies about 4°C across the day. Temperatures below this homeothermic point lead to increased energy expenditures, whereas temperatures above lead to a rise in body temperature [15].

For a naked human being the thermoneutral zone is similar to that of mice and ranges between 28°C and 29°C [26]. But as soon as the human body is covered with light clothing (e.g. long sleeved shirt or blouse and light trousers) this range drops down to 23°C - 25°C [27] or to 15°C - 25° with regular clothing (e.g. a business suit) [26]. Offering mice bedding and nesting material for insulation could have a comparable effect as clothing in humans. Thus, mice can adapt to different ambient temperatures, given that sufficient bedding and nesting material is available. Moreover, they are able to adjust their body core temperature depending on activity and environmental conditions and are even able to survive ambient temperatures from -10°C to 32°C [28]. Interestingly, this characteristic seems to be dependent on sex, strain, age or an interaction of these variables. For example, when kept at ambient temperatures between 20°C and 30°C, 6 months old C57BL/6 females showed a subcutaneous temperature difference of 0.5°C [24]. In contrast, 2 months old CD1 males kept in this temperature range showed a 2°C difference [13], and no difference in body temperature was found in 6 weeks old BALB/c females between 20°C and 30°C ambient temperature [11]. Even between phases of activity and inactivity mouse body temperature differed in about 1°C [24,29-32]. And at 20°C, mouse body temperature was not influenced by the presence or absence of nesting material, only food consumption was increased in the absence of nesting material [20]. Age [33] and strain [34] can influence experimental data that are collected at homeothermic (30°C) or common facility temperatures (20°C).

However, the question, which temperature mice prefer in regard of their wellbeing, is still open. Tumor bearing mice, i.e. morbid animals, preferred higher temperatures, because their thermoregulation is potentially already defective [35]. In preference tests healthy mice spent more time in warmer surroundings when they were inactive, i.e. slept or rested, or when solely cage bedding was available [16]. If, however, nesting material was offered and mice had the possibility to carry it over into cages with different ambient temperatures they allocated it in cooler cages and used it for nest building to insulate themselves while resting [17]. However, even if nesting material was provided a preference for a warmer environment of adult female mice was observed especially in the inactive phase, compared to male mice of the same age [36]. Possible effects of ambient temperatures on animal welfare have been addressed [29,30,32] and reproductive parameters like birth rate, weaning rate and embryo quality were investigated in relation to this environmental factor in mice [21,22,37,38]. Also, increased sleeping apneas [39] and behavioral changes, such as increased male aggression [40] were reported for mice in studies with higher ambient temperature.

Toth and coworkers [32] were the first to investigate the impact of ambient temperatures on animal welfare by measuring fecal corticosterone metabolites (FCMs) of mice kept at different room temperatures. Measuring FCMs is a proven non-invasive method to evaluate the animals’ stress hormone levels [41-44]. In the above mentioned study no difference in FCM concentration was found in adult C57BL/6J female mice when maintained at ambient temperatures of 22°C, 26°C or 30°C, but it must be noted that their FCM method was not validated [32,44].

Unfortunately, there are no studies to our knowledge, regarding the optimal ambient temperature for the wellbeing of lactating mice. Lactation is a highly demanding metabolic process [45,46] accompanied by considerable metabolic heat production as a by-product. Knowing the optimal ambient temperature of lactating mice would be highly valuable to optimize animal keeping and conditions in breeding colonies.

In this study we therefore investigated the impact of different ambient temperatures (20°C, 25°C, and 30°C) on the reproductive performance and wellbeing of female inbred (C57BL/6N) and hybrid (B6D2F1) mice during lactation compared to non-pregnant controls. We measured glucocorticoid metabolite levels in feces, animal food consumption, amount of voided feces and individual body weight. The reproductive performance was assessed by comparing the number of implantation sites, the number of born and weaned offspring, as well as adult and pup weight. In addition, we measured offspring tail length at weaning.

## Materials and Methods

### Animals and husbandry conditions

A total of 30 male C57BL/6N (referred to as B6) and 30 male B6D2F1 (referred to as F1) at the age of 8 weeks and 60 female B6 and 60 female F1 at the age of 6 weeks were purchased from Janvier Laboratories, Laval, France. Mice were specific pathogen free (SPF) according to FELASA recommendations and maintained in a barrier rodent facility. Groups of 3 to 4 females and single males were housed 2 weeks in type II Macrolon® cages for acclimatization. The cages were lined with bedding (Lignocel® Select, Rettenmaier KG, Austria) and enriched with nesting material (Arbocel® Crinklets natural, Rettenmaier KG, Austria; PurZellin, Paul Hartmann GesmbH, Austria) (photoperiod 12L:12D). Food (V1534 for males, non-pregnant females and females without pups, V1124 for pregnant females and females with pups, Ssniff Spezialdiaeten GmbH, Germany) and tap water were available *ad libitum*. Experimental procedures were discussed and approved by the institutional ethics and welfare committee and granted by the national authority according to §§ 26ff. of the Animal Experiments Act, Tierversuchsgesetz 2012 – TVG 2012 under license number BMBWF-68-205/0162-V/3b/2019.

### Experimental temperature groups

At the beginning all animals were housed at 20°C cage temperature under standard housing conditions as described above. To induce pregnancy in experimental mice females were mated bigam with a male of the same strain and checked daily for vaginal plugs. Every day, plug positive females were re-housed separated by strain in groups of 3 to 4. Within 4 days of permanent mating 37 females per strain were plug positive. These females were randomly assigned (12/12/13) to one of the temperature groups (30°/25°/20°C). In addition, 8 B6 and 8 F1 plug negative or non-mated females, and 8 B6 and 8 F1 males of the same age were used as controls for each temperature group. Seven days after the detection of a vaginal plug the group that was assigned to a 30°C cage temperature was transferred into an identical room next door with 25°C cage temperature for stepwise adaptation. After seven days, this group was finally re-located to an identical room next door with 30°C for the last week of pregnancy, birth and lactation. The second group was transferred to 25°C room 14 days after plug detection. The third group stayed in the room with 20°C cage temperature from the beginning and remained there until the end of the experiment (Fig 1). We expected pup births about 20 days after plug detection. Consequently, one week before the expected birth date all experimental and control animals were in rooms with their assigned cage temperature. Because birth took place between 18 to 21 days after plug detection the exact number of days under increased ambient temperatures before parturition differed slightly between animals of the respective temperature groups.

**Fig 1.**
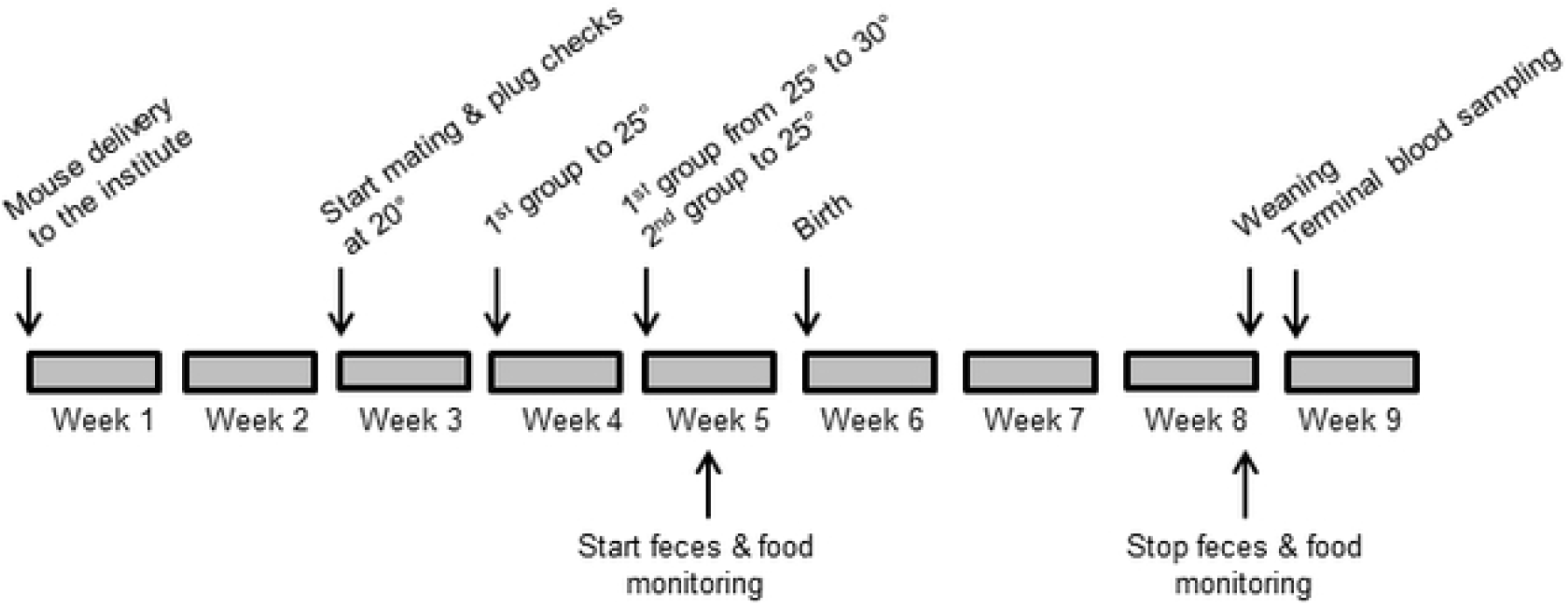
Experimental time schedule. Schematic description of the experimental manipulations and sample collections performed throughout the experiment.

## Experimental measurements

Cage temperature was measured with five temperature loggers per room (DS1921G, Thermochron, OnSolution Pty Ltd, NSW, Australia) deposited in 5 cages on different rack levels. Measurements were recorded every two hours. Humidity was recorded two times a day (at weekends only once) with standard hygrometers at 3 different positions in the room. We monitored pregnancies, and recorded the day of birth, the number of pups per litter at birth and at weaning. Over a period of 4 weeks, i.e. from last week of pregnancy until weaning, we measured animal food consumption once a week for 24 h. Therefore, we took the weight of the food in the hopper at the beginning and at the end of the 24 h period without spillage correction. Tail length of pups was measured on day 21 post birth with a digital caliper. Body weight of adults was measured at weaning using an electronic scale. For male and female controls the day of the first weaning in the experimental groups was used as reference. Individual pup weight of all litters was taken on the same day at a pup age between 16 to 21 days to assess intra-litter variation. For assessment of inter-litter-variation the whole litter weight was taken at weaning (d 20) and mean body mass was calculated by dividing the whole litter weight by the number of pups.

### Implantation sites

In order to evaluate the number of born pups in relation to the number of implanted embryos we dissected the uteri of breeding females *post mortem* at the end of the study. We opened the uterine horns with scissors and stained the implantation sites with a few drops of 10% ammonia solution [47]. After a few minutes of reaction implantation sites, visible as dark spots, were counted.

### Analysis of fecal corticosterone metabolites and plasma corticosterone

We sampled feces daily starting at the same time to determine fecal corticosterone metabolites (FCMs) excreted during activity phase over night in all mice. We started sample collection a few days before females gave birth and continued until weaning of the pups. Sample collection for controls occurred at the same dates. Due to the high number of samples only two per mouse and week were analysed. The first two time points were 1-3 days prior to birth (because of differing birth dates). The third sample time point was for mothers on the day of birth and for corresponding controls at the same day. Sample time points 4-9 followed in 3-4 days intervalls. The last time point was the day of weaning (Fig 1).

For sample collection, mice were put individually into clean pipette boxes for 15 minutes and fresh feces were collected. If the amount of voided feces during this time period was insufficient for analysis, respective mice were put into clean type III cages without bedding and the collection interval was prolonged for up to 30 minutes. Samples were stored at -20°C and FCMs were determined according to a routinely used protocol. Briefly, dried and homogenised feces were weighted and mixed with 80% methanol, centrifuged, the supernatant was diluted and an aliquot was analysed in a well-established and validated 5α-pregnane-3β,11β,21-triol-20-one enzyme immunoassay (EIA) [48,49].

Additionally, once a week a 24 h sample collection was performed. Therefore, animals were transferred to a fresh cage and after 24 h bedding and feces were collected and frozen. As voided feces were mixed with the fresh bedding we sorted the fecal pellets later by hand before weighing. The total amounts of excreted feces within 24 h was recorded in mice between temperature groups to be able to account for differences in food consumption and of droppings, respectively. If mice consume less food and secrete fewer droppings, this might lead to increased concentrations of FCMs per gram feces and *vice versa*.

After weaning and for controls at an equivalent time point all mice were sacrificed and blood was collected by heart puncture. Serum was prepared and analysed for blood corticosterone. Plasma samples were extracted with diethyl-ether and analysed with a previously described corticosterone EIA [50].

### Statistical procedures

Statistical analyses were performed with IBM SPSS Statistics 24.

To assess how cage temperature affected female reproduction we performed different models. First, we run a Generalized Linear Model (GLM) with a binomial distribution where we included the incidence of pregnancies as the dependent variable and we run a GLM with a poisson distribution, where we included the number of implantation sites, litter size at birth and at weaning as dependent variables. Finally, we performed Linear Models (LM), where we included litter weight at weaning, female body mass at weaning, mean pup body mass and pup tail lengths as dependent variables. Mouse strain and cage temperature were always included as fixed factors to all models and Least Significant Difference (LSD) Test was applied as post-hoc test to assess differences between temperature groups. We further tested whether the variation in individual pup body mass (SDs) within litters differed depending on their cage temperature with a Kruskal Wallis Test.

To assess how the experimental manipulations affected FCM levels, food consumption and feces production over the course of the experiment, we performed repeated measures ANOVAs. We included individual FCM levels, the calculated amount of daily food consumption, and the repeatedly recorded daily feces production as dependent variables, cage temperature, strain, animal sex and female breeding status as fixed factors. To assess differences within groups Least Significant Difference (LSD) Test was applied as post-hoc test. Finally, we also assessed plasma corticosterone levels with a LM where we included cage temperature and mouse strain as fixed factors. We tested in all models if model assumptions were fulfilled and transformed data if necessary.

## Results

### Cage temperature and room humidity

Experimentally intended cage temperatures were constantly maintained. Relative humidity decreased with increasing ambient temperatures. At 30°C air temperature humidity was comparatively more fluctuating, but at all times within the range of 30% to 50%.

### Reproductive parameters

Out of 74 females with a mating plug and additional two females without a plug, 54 (71.1%) became pregnant and 22 plugged females (28.9%) did not show any signs of gestation. Pregnancy rates were not affected by cage temperature (χ^2^=4.24, p=0.120), but were significantly higher in F1 compared to B6 females (χ^2^=11.90, p=0.001; Table 1).

**Table 1.**
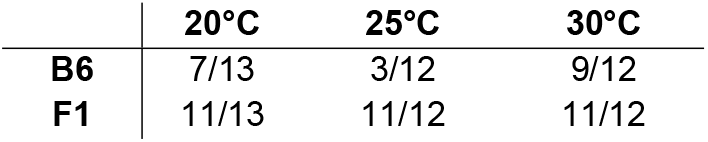
Number of parturient B6 and F1 females per plug positive females that were kept at 20°C, 25°C and 30°C.

|  | 20°C | 25°C | 30°C |
| --- | --- | --- | --- |
| **B6** | 7/13 | 3/12 | 9/12 |
| **F1** | 11/13 | 11/12 | 11/12 |

Females gave birth to an average of 7.5 pups per litter and litter size at birth did not differ between cage temperature (GLM: χ^2^=0.29, p=0.863) or female strain (GLM: χ^2^=1.63, p=0.202). Similarly, the number of female implantation sites (mean: 8.2) did not differ between cage temperature (GLM: χ^2^=0.09, p=0.957) or female strain (GLM: χ^2^=0.16, p=0.694).

We found that cage temperature had a significant effect on the number of pups weaned (GLM: χ^2^=7.19, p=0.027; Fig 2), and females kept at 30°C weaned fewer pups compared to females kept at either 20°C (p=0.042) or 25°C (p=0.002). No difference was found in the number of pups weaned in females kept at 20°C compared to 25°C (p=0.197). Also, F1 females weaned significantly more pups compared to B6 females (GLM: χ^2^=14.8, p<0.001; Fig 2). The number of litters corresponds to the number of females giving birth (Table 1).

**Fig 2.**
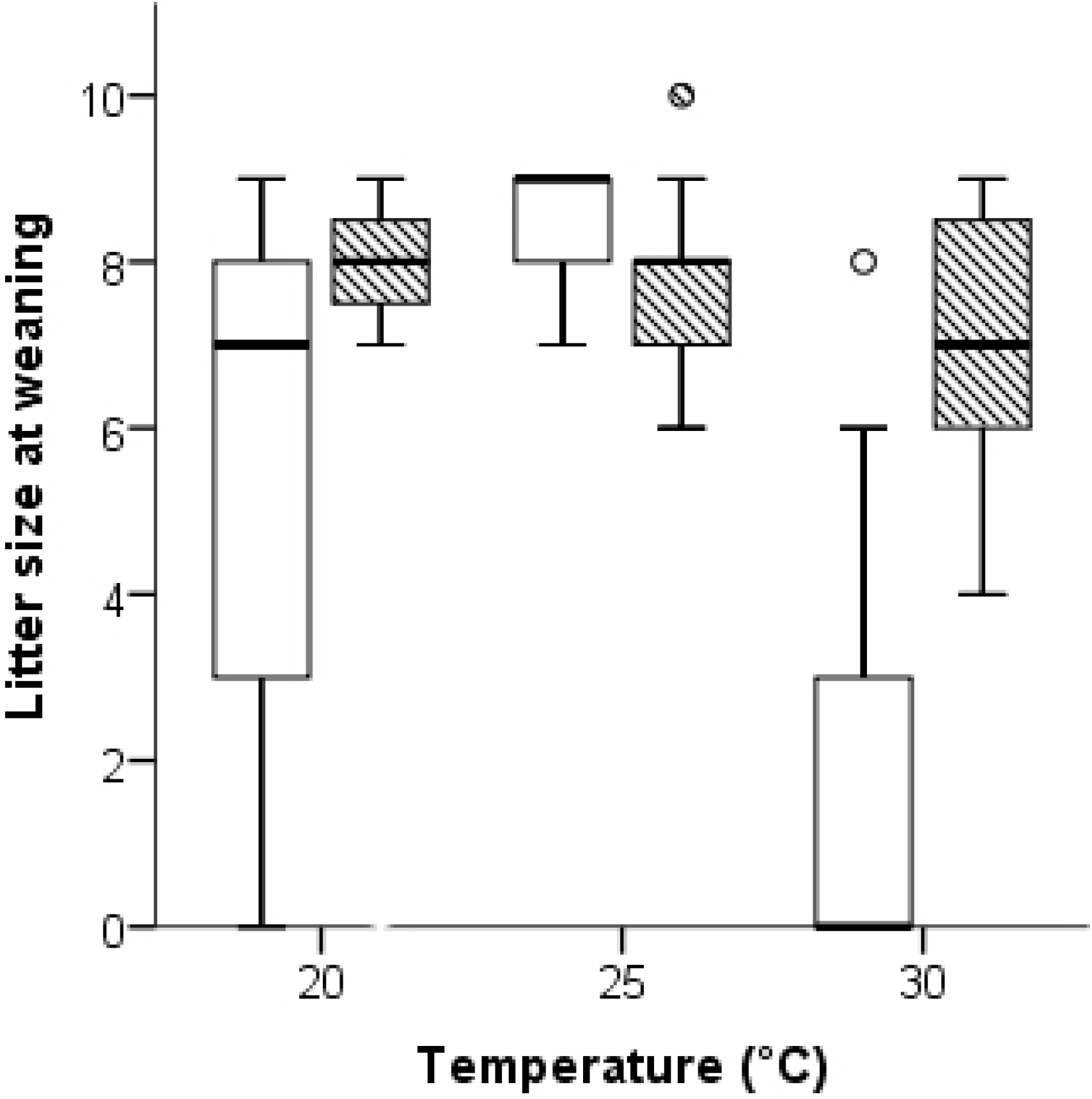
Boxplot of litter size at weaning in B6 (white boxes) and F1 hybrid (striped boxes) females kept at 20°C, 25°C and 30°C. Dot = mild outlier (Q1-1.5*IQ, or Q3+1.5*IQ).

### Weight and tail length

Similarly to litter size at weaning, we also observed that litter weight at weaning was significantly affected by cage temperature (F=17.71, p<0.001; Fig 3).

**Fig 3.**
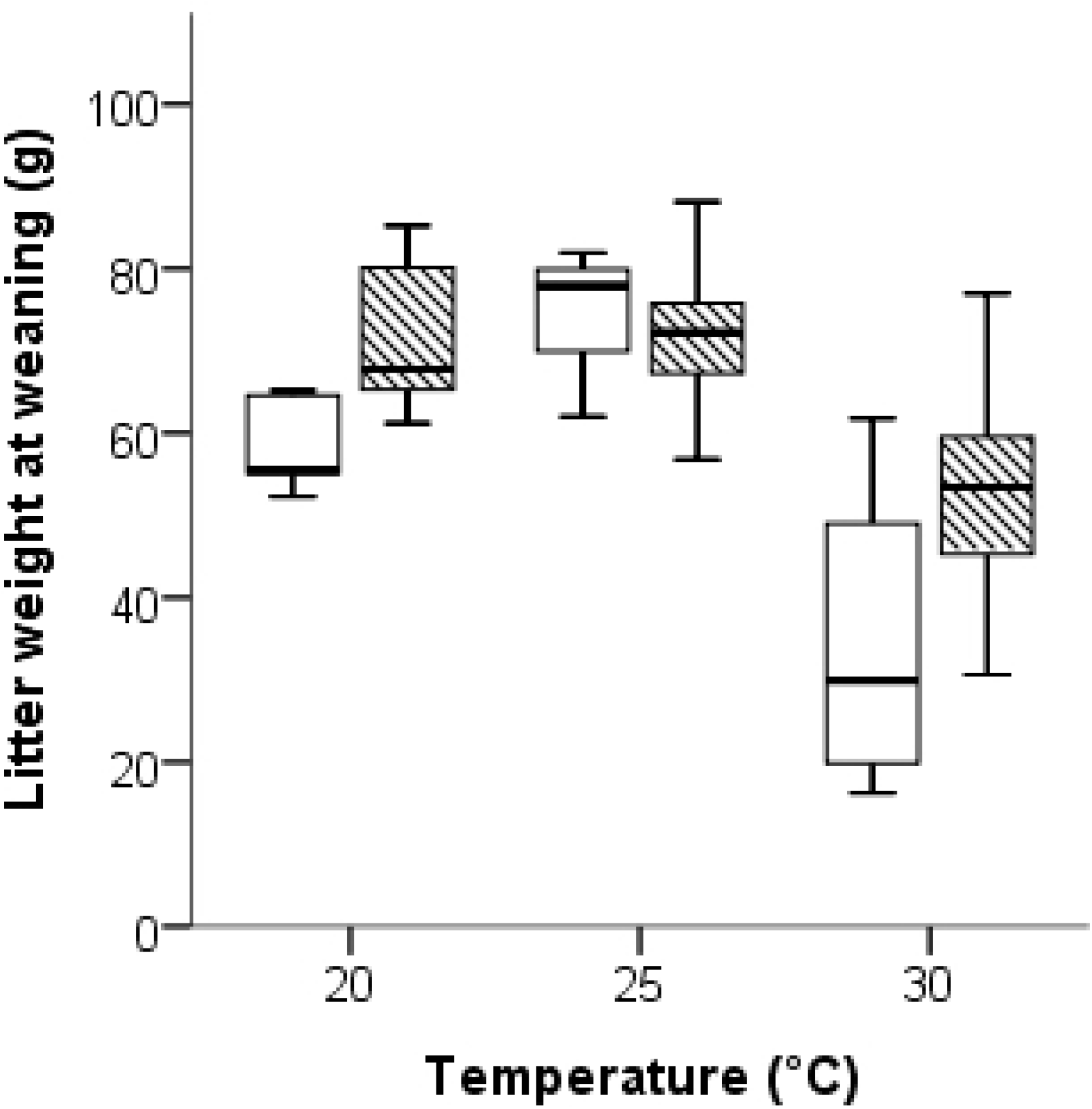
Boxplot of litter weight at weaning in B6 (white boxes) and F1 hybrid (striped boxes) females kept at 20°C, 25°C and 30°C.

Females kept at 30°C showed significantly lower litter weaning weights compared to females kept at 25°C (p<0.001) or 20°C (p<0.001). No difference in litter weaning weight was detected between females kept at 25°C or 20°C (p=0.218). Also, F1 females weaned significantly heavier litters compared to B6 females (F=7.94, p=0.007; Fig 3), though F1 mothers were significantly lighter than B6 mothers (F=8.88, p=0.005; Fig 4). Female body mass was also affected by cage temperature (F=70.64, p<0.001; Fig 4) and significantly declined with increasing temperatures (all post-hoc tests p≤0.011; see Supplement Information Fig S1).

**Fig 4.**
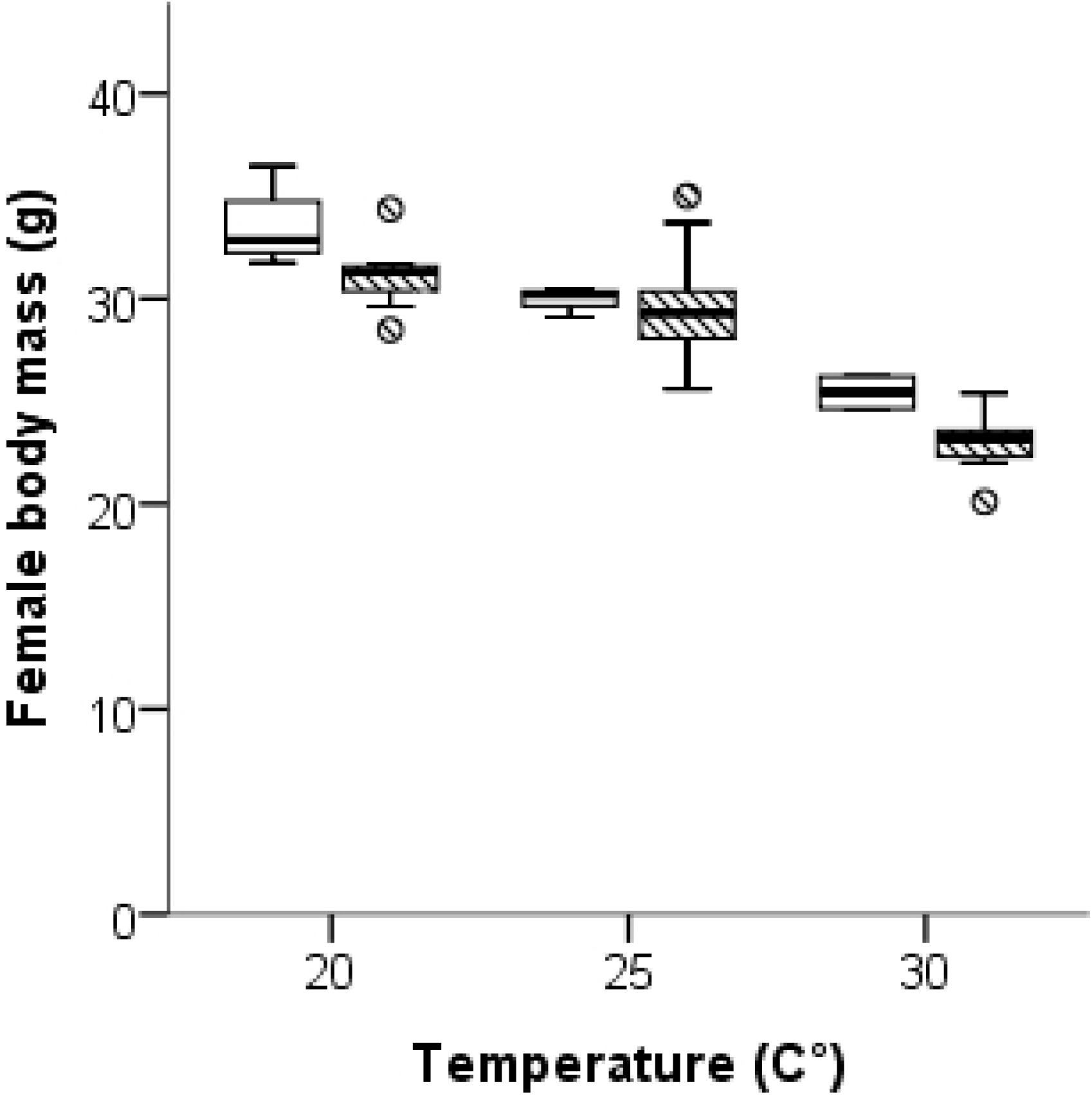
Boxplot of female body mass at weaning in B6 (white boxes) and F1 hybrid (striped boxes) females kept at 20°C, 25°C and 30°C. Only females that weaned pups are included in the graph. Dot = mild outlier (Q1-1.5*IQ, or Q3+1.5*IQ), asterisk = extreme outlier (Q1-3*IQ, or Q3+3*IQ).

Mean pup body mass also differed significantly between cage temperatures (F=13.39, p<0.001; Fig 5) and was highest in the 25°C group, followed by the 20°C group and was lowest in the 30°C group (all post-hoc tests p≤0.025). We did not detect any strain specific differences in mean pup body mass (F=3.34, p=0.075; Fig 5), and we did not notice any differences in the within litter variation in body mass depending on cage temperature (Kruskall Wallis Test: p=0.389).

**Fig 5.**
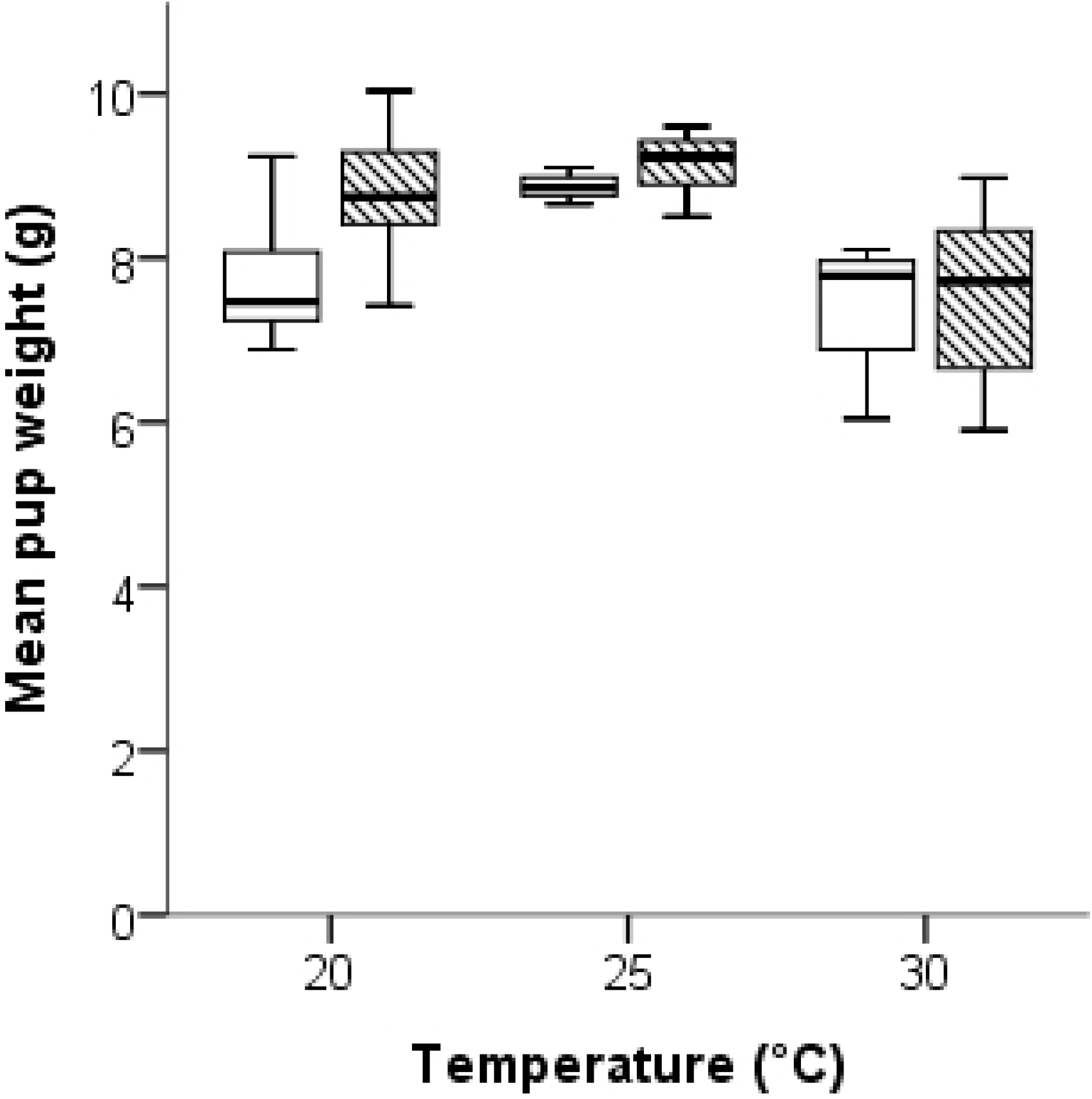
Boxplot of mean pup weight at weaning in B6 (white boxes) and F1 hybrid (striped boxes) females kept at 20°C, 25°C and 30°C. Asterisk = extreme outlier (Q1-3*IQ, or Q3+3*IQ).

Finally, we found that the mean tail length of litters was affected by both, female strain (F=31.92, p<0.001; Fig 6) and cage temperature (F=67.32, p<0.001; Fig 6). Pups of F1 females had on average longer tails compared to offspring of B6 females and pups from mothers of both strains kept at 20°C had significantly shorter tails compared to pups from mothers kept at either 25°C (p<0.001) or 30°C (p<0.001). No difference in pup tail length was found between 25°C and 30°C (p=0.356).

**Fig 6.**
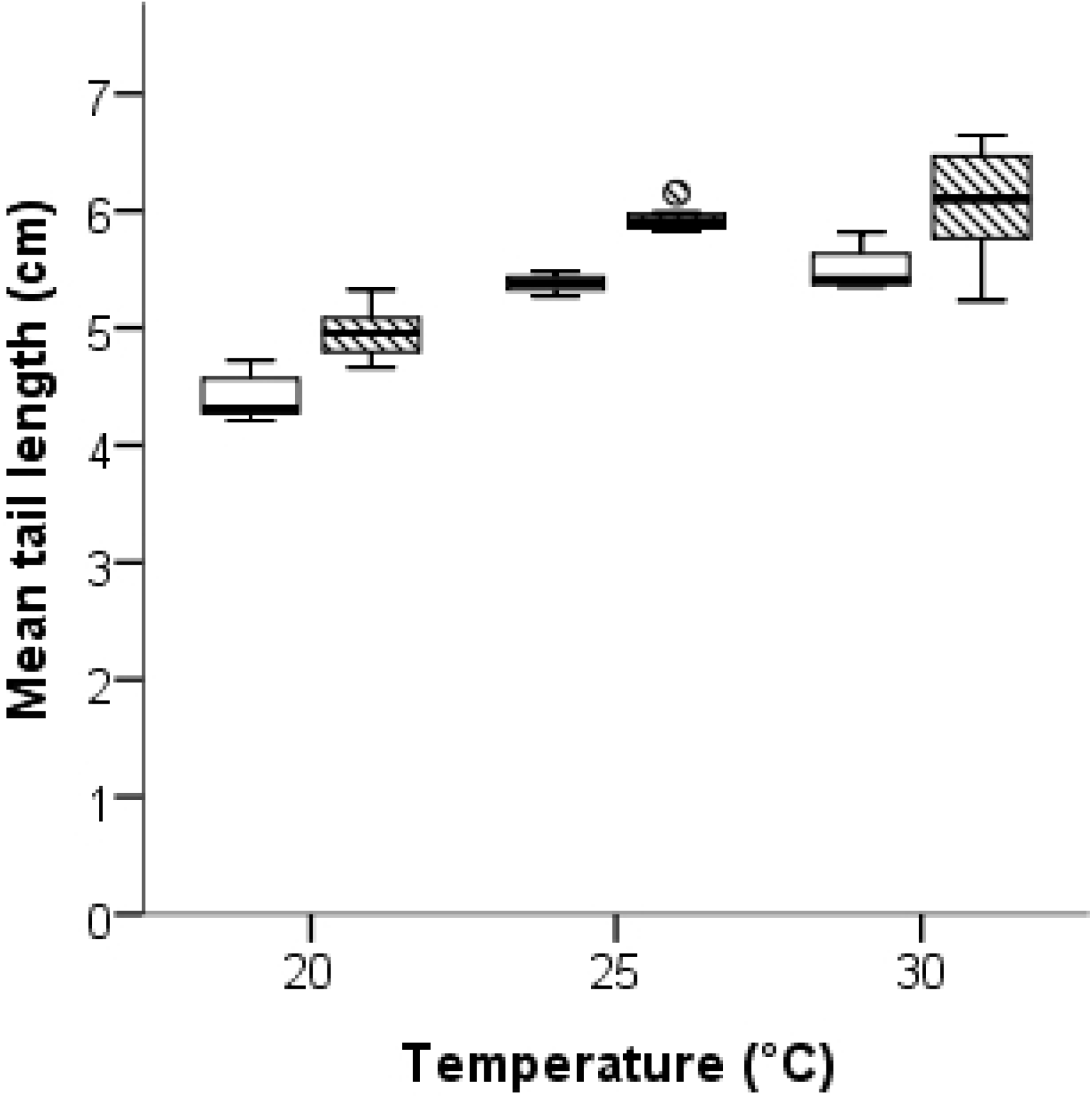
Boxplot of mean tail length in pups weaned from B6 (white boxes) and F1 hybrid (striped boxes) females kept at 20°C, 25°C and 30°C. Dot = mild outlier (Q1-1.5*IQ, or Q3+1.5*IQ).

### Food consumption and amount of feces

When investigating animal food consumption, we found that F1 hybrid mice consumed on average significantly more food per day compared to B6 mice (F=21.12, p<0.001; Fig 7B). Also, daily food intake was affected by cage temperature (F=27.58, p<0.001; Fig 7A) and was reduced significantly with rising cage temperatures (all post-hoc tests: p≤0.002). In addition, food intake also varied between mice depending on their sex and breeding status (F=49.56, p<0.001; Fig 7C). Experimental (breeding) females consumed significantly more food compared to mice from the control groups (p<0.001). No difference was found between male and female control mice (p=0.535). In line with the higher food consumption, F1 hybrids produced significantly more feces per day than B6 mice (F=19.48, p<0.001; Fig 8B). Moreover, feces production significantly decreased in parallel to food consumption with rising ambient temperatures (F=29.72, p<0.001; Fig 8A; all post-hoc tests: p<0.001). Finally, daily feces production varied between mice depending on their sex and breeding status (F=41.76, p<0.001; Fig 8C) and breeding females produced significantly more feces compared to mice from the control groups (p<0.001). Again, no difference was seen between female and male control mice (p=0.539).

**Fig 7.**
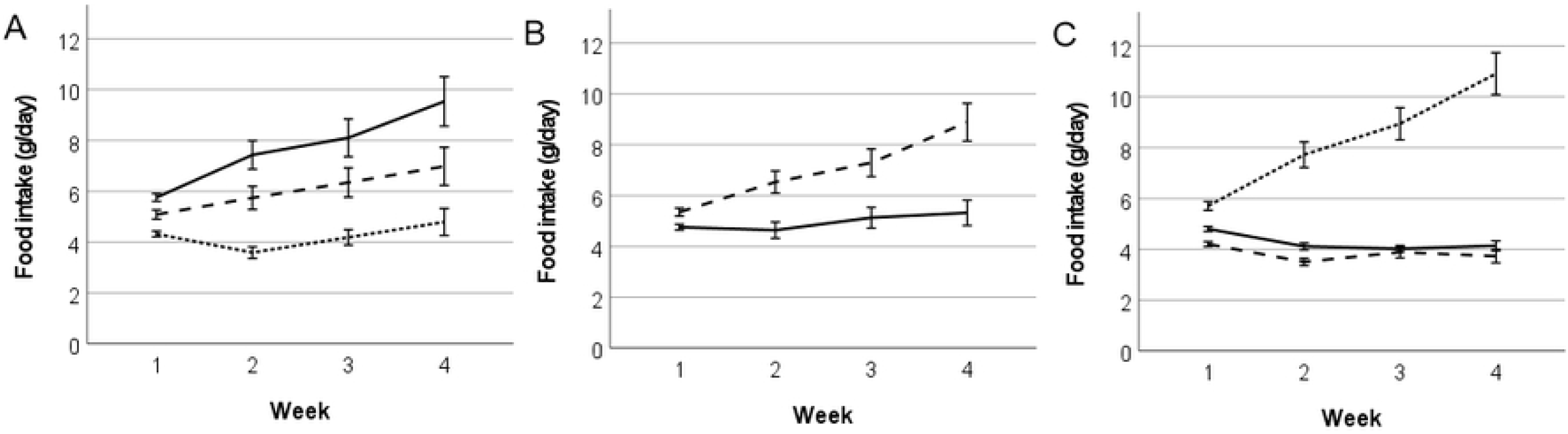
Mean (±SE) animal food consumption over a period of 4 weeks in male, non-reproducing female and reproducing female B6 and F1 mice kept at 20°C, 25°C and 30°C. (A) Food consumption in mice kept at 20°C (solid line), 25°C (dashed line) and 30°C (dotted line). (B) Food consumption in B6 (solid line) and F1 (dashed line) mice. (C) Food consumption in male (solid line), non-reproducing female (dashed line) and reproducing female (dotted line) mice.

**Fig 8.**
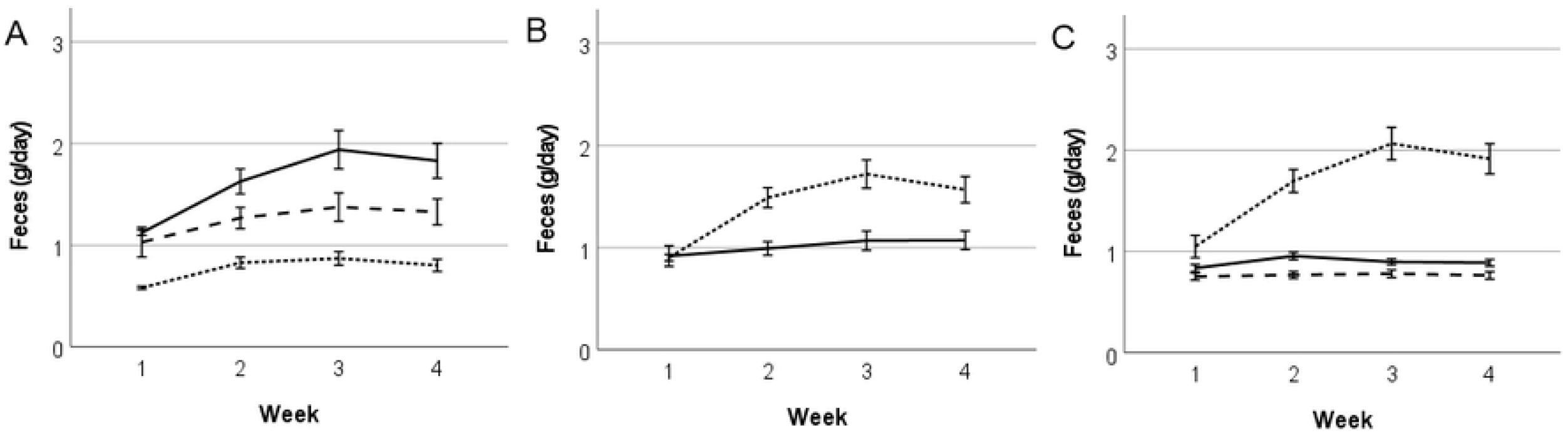
Mean (±SE) animal feces production per 24 h over 4 weeks in male, non-reproducing female and reproducing female B6 and F1 mice kept at 20°C, 25°C and 30°C. (A) Feces production in mice kept at 20°C (solid line), 25°C (dashed line) and 30°C (dotted line). (B) Feces production in B6 (solid line) and F1 (dashed) mice. (C) Feces production in male (solid line), non-reproducing female (dashed line) and reproducing female (dotted line) mice.

### Fecal corticosterone metabolites (FCMs) and plasma corticosterone

FCM levels differed significantly between mouse strains (F=42.78, p<0.001; Fig 9B), as F1 mice showed constantly higher values compared to B6 mice. In addition, FCM levels differed significantly between mice depending on their sex and breeding status (F=305.86, p<0.001; Fig 9C): Breeding females showed significantly higher FCM levels compared to both, control females and males (p<0.001) and control females showed significantly higher FCM levels compared to control males (p<0.001). Interestingly, breeding females showed peak values in FCM levels at the time of birth and at weaning of their offspring. However, FCM levels did not differ between mice depending on their cage temperature (F=0.71, p=0.493; Fig 9A).

**Fig 9.**
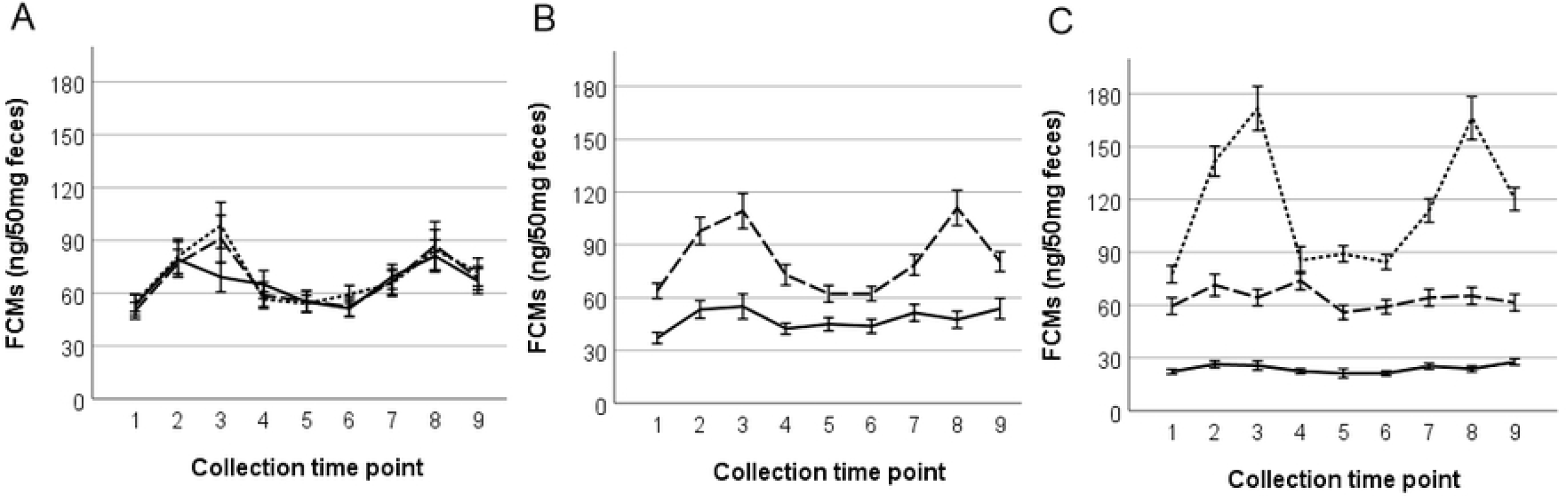
Mean (±SE) FCMs over time in male, non-reproducing female and reproducing female B6 and F1 mice kept at 20°C, 25°C and 30°C. FCMs= Fecal corticosterone metabolites. (A) FCM levels in mice kept at 20°C (solid line), 25°C (dashed line) and 30°C (dotted line). (B) FCM levels in B6 (solid line) and F1 (dashed line) mice. (C) FCM levels in male (solid line), non-reproducing female (dashed line) and reproducing female (dotted line) mice. Peak values were observed at birth (time point 3) and shortly before weaning (time point 8).

Finally, we observed that plasma corticosterone levels at the end of the experiment confirmed the findings of the FCM analysis and did not show any difference between strains (F=0.0, p=0.997) or temperature groups (F=2.89, p=0.059; data not shown).

## Discussion

### Reproduction

In our study we investigated the effect of different housing temperatures (20°C, 25°C, 30°C) on breeding performance and stress levels in female C57BL/6N (B6) inbred and D2B6F1 (F1) hybrid mice.

As expected from hybrid vigor, we found that pregnancy rates after a four days mating period were significantly higher in F1 compared to B6 females. Neither pregnancy rate nor litter size at birth differed between experimental temperature groups, confirming that there was no bias in reproductive traits before the treatment started. This result is not surprising, because mating and the beginning of the pregnancy took place at 20°C for all experimental females. In line with this, cage temperature and strain had no effect on the number of implantation sites. The low number of 3 pregnant B6 females out of 12 pluged after mating in the 25°C group seems to be merely an unfortunate divergence.

All measured postnatal parameters like litter size and mean pup body mass at weaning were significantly affected by cage temperature and reached their poorest outcome in females kept at 30°C. The low number of pregnant B6 females in the 25°C group was considered in the statistical tests. As expected, the proportion of weaned pups was higher in F1 compared to B6 females. Interestingly, the impact of a 30°C cage temperature on reproduction was more pronounced in B6 females, suggesting an increased sensitivity of this inbred strain to high ambient temperatures, whereas hybrids seemed to better tolerate heat. The observerd impact of higher ambient temperatures on reproduction is similar to results from Yamauchi and coworkers [37], who described decreased litter sizes and increased pup losses in ICR outbred mice kept at temperatures from 26°C to 32°C. In another study with SWISS mice, milk production at 33 °C was only 18% of that at 21 °C. This led to reductions in pup growth by 20% but only limited pup mortality (0.8%) was observed [51]. In contrast to our study with a heat exposure starting at the last third of pregnancy, Zhao and coworkers exposed the females and their litters only from day 6 postpartum to higher temperatures, whereas the pup losses in our study occurred only during the first 24 hours after birth. In rats kept at 33°C [52] and hamsters kept at 30°C [53,54] a negative temperature effect was also observed on reproductive parameters. In our study the best reproductive results were found when females were kept at 25°C, though there was hardly any significant difference between 20°C and 25°C. Interestingly, F1 females showed consistently better reproductive outcomes compared to B6 and over all temperature groups, indicating that these hybrid females are better able to cope especially with higher temperatures.

### Physiological and morphological changes

The cage temperature also influenced other physiological and morphological parameters like body weight of lactating mothers and tail length in pups. Females kept at 30°C were significantly lighter, compared to females at either 20°C or 25°C. The lower body weight at 30°C could be explained by the reduced food consumption in this group. In line with this, also mean pup body mass was significantly lower at 30°C compared to either 25°C or 20°C and is in accordance with other studies [55-57]. Pup body mass is directly related to female body mass since the development of the mammary gland and lactation is dependent on adequate food intake. Alternatively, and not mutally exclusive, pup body mass can further be affected by the impact of the ambient temperature on the lactating mother: According to the heat dissipation limit hypothesis, females cannot dissipate enough metabolic heat at higher ambient temperatures and therefore limit milk production, which results in reduced pup weight [58,59]. This hypothesis was critically discussed by Sadowska and coworkers [60]. Nevertheless, higher ambient temperatures lead to reduced mammary glands [61] and additionally to reduced energy, fat and total solids in the milk [62] resulting in reduced growth of sucklings. It was also shown that milk energy output and suckling time were lower at 30 °C independent from the litter size [63].

We further found that pups from mothers kept at either 25°C or 30°C had significantly longer tails compared to pups from mothers that were kept at 20°C. The finding of longer tails in mice reared at high temperatures was reported previously [16,64]. A recent paper challengend the general assumption that the hairless and rich vascularized tail of mice is an important structure for the dissipation of body heat [65]. However, the observed elongation of the tail at this early developmental stage could be interpreted as an increase of the relative importance of the tail in its function to get rid of body heat under conditions of so-called homeothermy. This is an extremely quick adaptation, which was certainly facilitated by the postnatal growth period. Tail elongation as a so-called warm adaption was also detectable in adult BALB/c females if juveniles from 5 weeks of age were henceforth permanently exposed to high ambient temperatures [15]. In addition, we also found that pups of hybrid females had on average longer tails than offspring of B6 females. The finding confirme the results of Harrison and coworkers (1959) [64]. Because mean pup body weight at weaning was similar in the elevated temperature group in both strains, the more distinct tail elongation of hybrids indicates that the heterozygous background of hybrid mice facilitates a faster and better adaptation to increasing ambient temperatures than the homozygous inbred strain.

### Glucocorticoids

FCM levels assessed from late pregnancy to weaning and plasma corticosterone levels at the end of the experiment did not differ between mice across cage temperature groups, suggesting that none of the chosen ambient temperatures was more or less stressful for the mice. Alternatively, mice might have perceived specific temperatures as stressful, but could have behaviorally adjusted to them, i.e. built a warm nest and spend more time in it at lower temperatures, or reduce their activity and try to cool at cage walls at higher temperatures. We did not permanently conduct observations to confirm behavioural adaptitions. However, we noted reduced nest building activity in the 30°C group (see Supplement Information Fig S2).

We found that hybrid mice showed constantly higher FCM levels compared to B6 mice. This is an interesting observation, because the detected plasma corticosterone levels of blood samples taken one day later did not show any difference between temperature groups or strains. Differences in FCM levels between strains are known from another study [40] and might be explained by genetic differences and not by differences in experienced stress levels, as both strains were treated identically. We found that FCM levels differed significantly between mice depending on their sex and breeding status. Sex differences in FCM levels are also well described [48,49] und our results confirm that males have generally lower values than females.

Not surprisingly, we further found a difference in FCM levels based on female reproductive status. Breeding females had significantly higher levels than control females. Interestingly, breeding females showed their peak values in FCM levels at the time of birth and in the third/last week of lactation. Similarly, a perinatal increase of FCM levels was also reported by Möstl and Palme [66].

It seems that birth itself, like in many other mammals, and the challenge between a decreasing milk supply at the end of the weaning period combined with an increasing food requirement in offspring is most stressful for reproducing females.

The question emerged whether more food intake and higher amounts of feces lead to lower FCM concentrations. Studies in cows [67] and rats [68] showed that increased food intake causes a higher metabolic rate, a higher glucocorticoid clearance rate, and therefore, more FCM excretion via feces. Interestingly, reproducing females, which consumed more food and produced more feces, still had higher FCM levels. Therefore, the FCM concentration in the feces is not dependent on the total amount of excreted feces and a correction in our study was not necessary.

## Conclusions

It is unquestionable that ambient temperature can have a major impact on mouse physiology, from heart rate and blood pressure [7] to tumor growth [35,69,70] and immunological parameters [69,70]. However, also other external factors such as humidity, microbiological status, light intensity, noise, nutrition, and others are known to have an impact [71-74].

Our results showed that neither a low (20°C) nor a high cage temperature (30°C) resulted in changed stress hormone levels in experimental animals. Unlike the statement about ›permanent cold stress‹ of other authors [2,15] a cage temperature of 20°C to 25°C was not connected to increased stress levels. Therefore, it may be concluded from our study that the ›cool‹ standard temperature in rodent facilities (21 +/-1°C) has most likely no negative effect on animal welfare, as long as nest building material is provided. In contrast, high ambient temperatures can reduce the number of surviving pups and induce specific physiological adaptations (increased tail length, reduced body weight) when exceeding a certain level.

Furthermore, room temperatures of around 30°C could be challenging for employees working tightly dressed in a mouse facility [38,75]. In consideration of our findings, we definitely cannot recommend a homeothermic cage temperature of 30°C for breeding mice.

## Supporting information

**S1 Fig 1**. Examples of lactating B6 (a, c) and F1 (b, d) females in the third week at 20°C (a, b) and 30° C (c, d).

**S1 Fig 2**. Examples of cages with B6 (a, c) and F1 (b, d) pups in the third week at 20°C (a, b) and 30°C (c, d).

## Acknowledgements

The excellent technical assistance of A. Peham for animal work and N. Krotky, T. Bernthaler, D. Batkay, C. Winding-Zavadil, K. Slavnitsch and Edith Klobetz-Rassam for lab work is gratefully acknowledged.

## Notes

### Competing Interest Statement

The authors have declared no competing interest.

